# Conservative taxonomy and quality assessment of giant virus genomes with GVClass

**DOI:** 10.1101/2024.08.15.607234

**Authors:** Thomas M Pitot, Tomàš Brůna, Frederik Schulz

## Abstract

**Background:** Large double-stranded DNA viruses of the phylum Nucleocytoviricota (Giant viruses; GVs) include the largest known viruses, both in terms of capsid and genome size and are associated with a wide range of eukaryotic hosts. The ones able to infect protists and algae have been shown to be the dominant orders of GVs in the environmental samples. These viruses encode for genes that may have significantly impacted biogeochemical cycling and host genome evolution. While GVs are frequently found in environmental sequence data, their large and complex genomes, composed of genes acquired from various cellular lineages, pose challenges for their identification and taxonomic classification.

**Results:** We present GVClass, a tool that identifies giant viruses in sequence data and provides taxonomic assignments, and estimates for genome completeness and contamination. GVClass performs gene calling optimized for giant viruses and utilizes a conservative approach based on consensus single protein phylogenies for robust taxonomic assignments. The genes used for classification represent highly conserved giant virus orthologous groups and low copy number cellular and viral panorthologs. In our benchmarking, GVClass demonstrated high quality and accurate taxonomic assignment of giant virus sequences. GVClass showed high to very high precision, with over 90% of tested instances correctly predicted at the genus level and near-perfect prediction (>99%) at higher taxonomic ranks (family, order, class).

**Conclusion:** In the light of rapidly increasing amounts of sequence data and associated metagenome-assembled genomes, GVClass provides a conservative approach to identify, classify and quality-check giant virus genomes, which with other methods often remained unassigned or misclassified using other methods. GVClass has already been used through viral meta-analysis and to benchmark the viral sequences detection pipeline geNomad. The standalone version is freely available and it has been integrated in the Integrated Microbial Genomes / Virus database (IMG/VR), offering the opportunity to upload user data for giant virus classification.

## Introduction

Giant viruses (GVs) of the viral phylum *Nucleocytoviricota* infect a wide range of eukaryotic hosts, exhibit a broad spectrum of morphologies, capsid sizes, and genome lengths, and are currently classified into five distinct orders: *Algalvirales, Asfuvirales, Chitovirales, Imitevirales*, and *Pimascovirales* (Schulz et al, 2022). Their double-stranded DNA genomes range from under 100 kb to over 2.7 Mb, encoding hundreds to thousands of genes, often acquired through dynamic gene exchanges with various cellular and viral lineages (Filée & Chandler, 2010, La Scola et al., 2003, Schulz et al., 2017, 2022). Among them, the presence of genes associated with diverse microbial metabolic pathways, including photosynthesis, the tricarboxylic acid (TCA) cycle and glycolysis, suggests that GVs may influence their host’s metabolic capabilities (Moniruzzaman et al, 2020, Schulz et al, 2020). With the increased availability of metagenomic sequencing data from diverse ecosystems, it has become possible to study the diversity and the distribution of giant virus genomes, greatly expanding our understanding of their genetic diversity, function, and role in shaping the ecosystems they inhabit.

Despite their complex genomes posing challenges to taxonomic classification, GVs encode several giant virus orthologous groups (GVOGs)(Aylward et al, 2021). A subset of these GVOGs serve as phylogenetic markers and are an integral to computational pipelines and tools for detecting and identifying GV in metagenome assembled sequences (contigs, bins) (e.g. Virsorter2, (Guo et al, 2021), geNomad, (Camargo et al, 2023), ViralRecall, (Aylward and Moniruzzaman, 2021) or TIGTOG, (Ah and Aylward, 2024)). Before the normalization of GVOGs usage, other tools were designed and based on metagenome reads alignment with giant virus open reading frames (ORFs) (Verneau et al, 2016, Kerepesi and Grolmusz, 2017). Recently, a set of 7 mainly vertically inherited GVOGs was used to establish a taxonomic framework for studying *Nucleocytoviricota* diversity (Aylward et al, 2021). The largest proportion of this framework is based on giant virus metagenome-assembled genomes (GVMAGs) (Moniruzzaman et al, 2020, Schulz et al, 2020).

GVMAGs were recovered from de novo assemblies and in some cases metagenomic binning is prone to errors, leading to fragmented, or mixed population genomes with potential cellular contamination (Schulz et al, 2020). Further, GVMAGs often contain only few conserved genes, making classification more challenging.

Here we present GVClass, a bioinformatic pipeline that assigns taxonomy to putative giant virus genomes down to species level. GVClass employs a conservative approach, relying on the consensus of single protein phylogenetic trees inferred from identified GVOGs in GV sequences after optimized gene calling. It also incorporates specific hallmark genes from the recently proposed *Mriyaviricetes* and *Mirusviricota* (Yutin et al, 2024, Gaïa et al, 2023), as well as cellular single-copy panorthologs (sets of genes that are orthologous across multiple species or lineages within a taxonomic group). Subsequently, GVClass estimates genome completeness and contamination of identified giant viruses and summarizes all results. Benchmarking *in silico* modified giant virus genomes showed the robustness of GVClass across different levels of completeness and taxonomic novelty. Taken together, GVClass is a fast, reliable and user-friendly tool to detect, classify and assess quality of giant virus genomes including those recovered from environmental sequence data as shown in Pitot et al, 2024.

## Results and discussion

### The GVClass framework

To identify and classify GVs, query sequences can be provided as either FASTA amino acid (faa) or nucleotide (fna) format input (Fig. 1). To benefit from the entire set of functionalities of GVClass, fna is the preferred input format, with a recommended minimum assembly size of 20kb.

**Fig 1:**
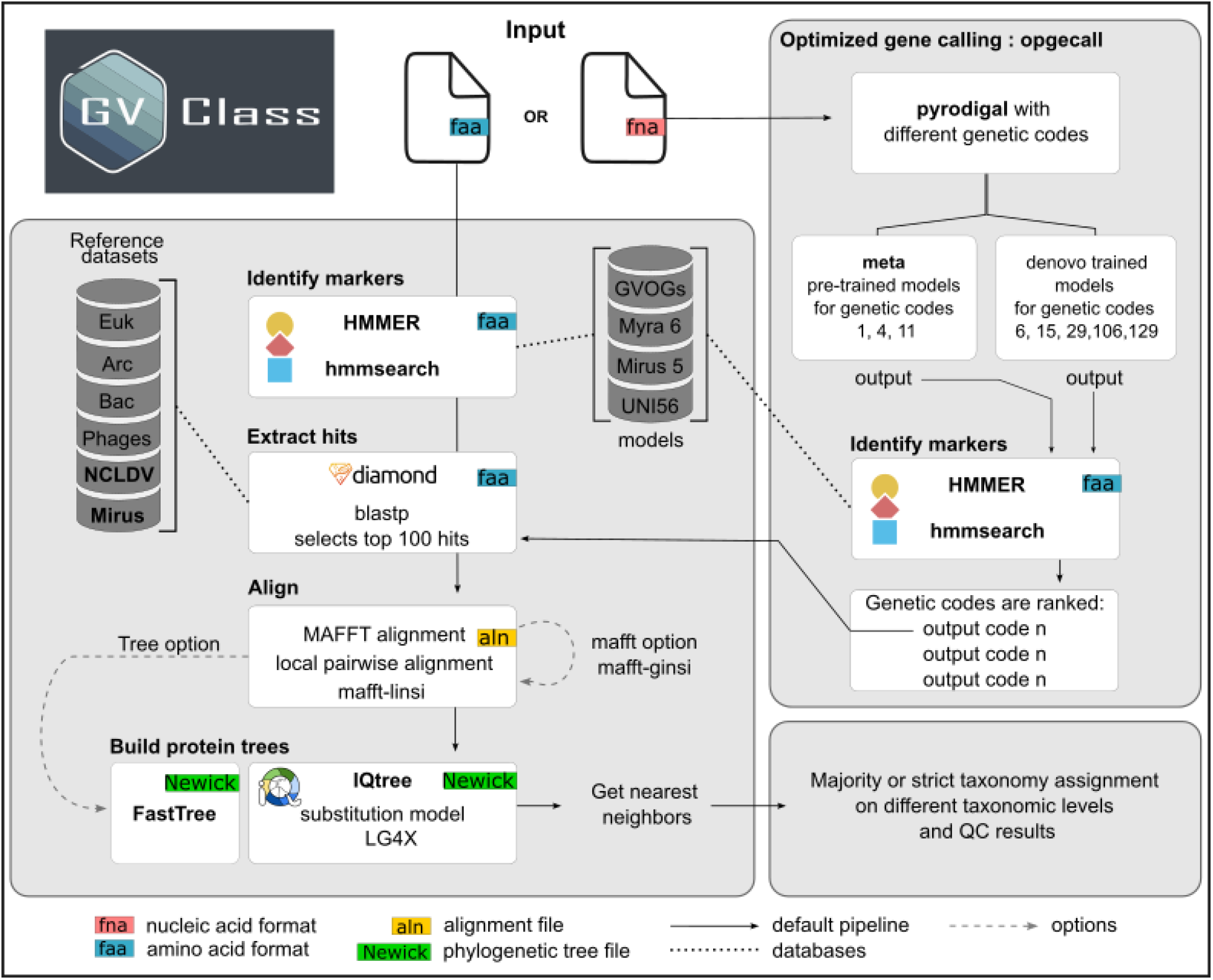
Overview of the GVClass framework. Schematic of the optimized gene calling (opgecall), taxonomic affiliation and quality assessment of giant virus sequences identified by GVClass

For fna input, nucleotide sequences undergo optimized gene calling (opgecall) using a modified version of pyrodigal (https://github.com/tomasbruna/pyrodigal) using different genetic codes: “meta” which employs pre-trained models for codes 1, 4, and 11, as well as *de novo* trained models for genetic codes 4, 6, 15, 29, 106 and 129. Codes 106 and 129 are our custom modifications to codes 6 and 29 where only one stop codon is re-assigned: code 106 re-assigns TAA to glutamine, and code 129 re-assigns TAG to tyrosine. Following the opgecall step, the outputs are assessed using hmmsearch (http://hmmer.org/v3.3.2) with a combined set of general HMMs and Nucleocytovirictoa order-level HMMs (Figure 1). Genetic codes are then ranked based on; 1) the highest number of complete profile hits (>60% of model coverage), 2) the average of bitscores corresponding to the best profile hits for each predicted protein. and 3) the coding density, which must exceed prodigal meta by 2% to select a *de novo* trained model. The highest ranked output is selected for subsequent analyses.

Hits with the general HMMs are extracted from translated proteins (faa), and each marker sequence is used as a query for a diamond blastp search against a custom reference database. This database contains pre-extracted homologs of the general HMMs from a representative set of bacterial and archaeal genomes from GTDB database (release 214) (Parks et al, 2022), EukProt (v3) (Richter et al, 2020) and IMG/VR (v4) (Camargo et al, 2022). The top 100 hits for each query are then extracted and aligned with MAFFT (v7.505) using local pairwise alignment (-linsi) by default. Alignments are then trimmed with trimAl (v1.4.1) (Capella-Gutiérrez et al, 2009) with the option -gt 0.1.

Next, single protein trees are constructed for each aligned set of proteins using IQ-TREE(v2.2.0.3) (Nguyen et al, 2015) with the LG4X substitution model (users can specify fastTree for slightly decreased runtime per tree). GVClass then analyzes phylogenetic trees to identify the nearest neighbors based on branch length of every query sequence, including paralogs. It employs functions to traverse the trees, find closest relatives, identify and characterize the nearest neighbors in provided reference datasets, and compile results.

To yield taxonomic assignment in the “strict” mode, all nearest neighbors (nn) in all phylogenetic trees must agree on the respective taxonomic level (species, genus, family, order, class, phylum, domain). In the “majority” mode (recommended), 50% of nn must agree for successful taxonomic classification. GVClass offers the option to build additional trees from a larger set of *Nucleocytovirictoa* order-level HMMs to increase taxonomic affiliation precision (fast_mode FALSE). However, as this leads to a much larger set of trees (up to 264) runtime of GVClass increases substantially.

For sequences predicted to be affiliated to the *Nuclectoviricota* phylum, the hit count of the order-level subset of the models (n=274) (see methods) is used to estimate lineage-specific genome completeness and contamination. For the latter, a lineage-specific duplication factor is calculated to estimate contamination. The genome completeness estimate in GVClass is based on the proportion of mainly single copy genes (duplication factor in published genomes <1.5) conserved in at least 50% of genomes of the respective *Nucleocytoviricota* order. In brief, the total number of hits is divided by the unique hits, taking into account the order-level conserved genes.

Finally, the GVClass pipeline summarizes all results in tabular output. The provided “gvclass_out_v1.0.0.tab” encompasses essential genome stats such as contig count, base pair length, GC content, gene count, coding percentage, and genetic code. Taxonomic classification is detailed, extending down to the species level whenever possible, for both “strict” consensus and a “majority” consensus. Moreover, the file offers detailed insights into specific genetic elements, including the unique and total counts of Nucleocytoviricota and Mirusviricota MCP (Major capsid protein) genes, GVOG4 and GVOG8 (a set of 4 and 8 GVOGs suitable for supermatrix-based species tree calculation), phage and universal cellular housekeeping genes, along with their respective duplication factors. As there is no universal rule for giant viruses(Claverie and Abergel, 2016), the provided wealth of information empowers users to discern intricate details about their query sequences.

Of particular significance are the duplication factor and completeness index provided at the predicted order level. A high duplication factor might suggest a sequence composed of multiple closely related giant viruses, while a low completeness index could indicate insufficient sequencing depth. Understanding these metrics aids users in interpreting their data accurately and making informed decisions regarding subsequent analyses.

### The GVClass benchmark

#### *Taxonomic classification with* GVClass

For benchmarking, we excluded self-hits and limited taxonomic classification to the Class, Order, Family and Genus ranks. We used a comprehensive set of published giant virus genomes (n = 1,109), which were fragmented and randomly reduced (in triplicate) to 75% 50% and 25% of their original length (Fig.2A).

**Fig 2:**
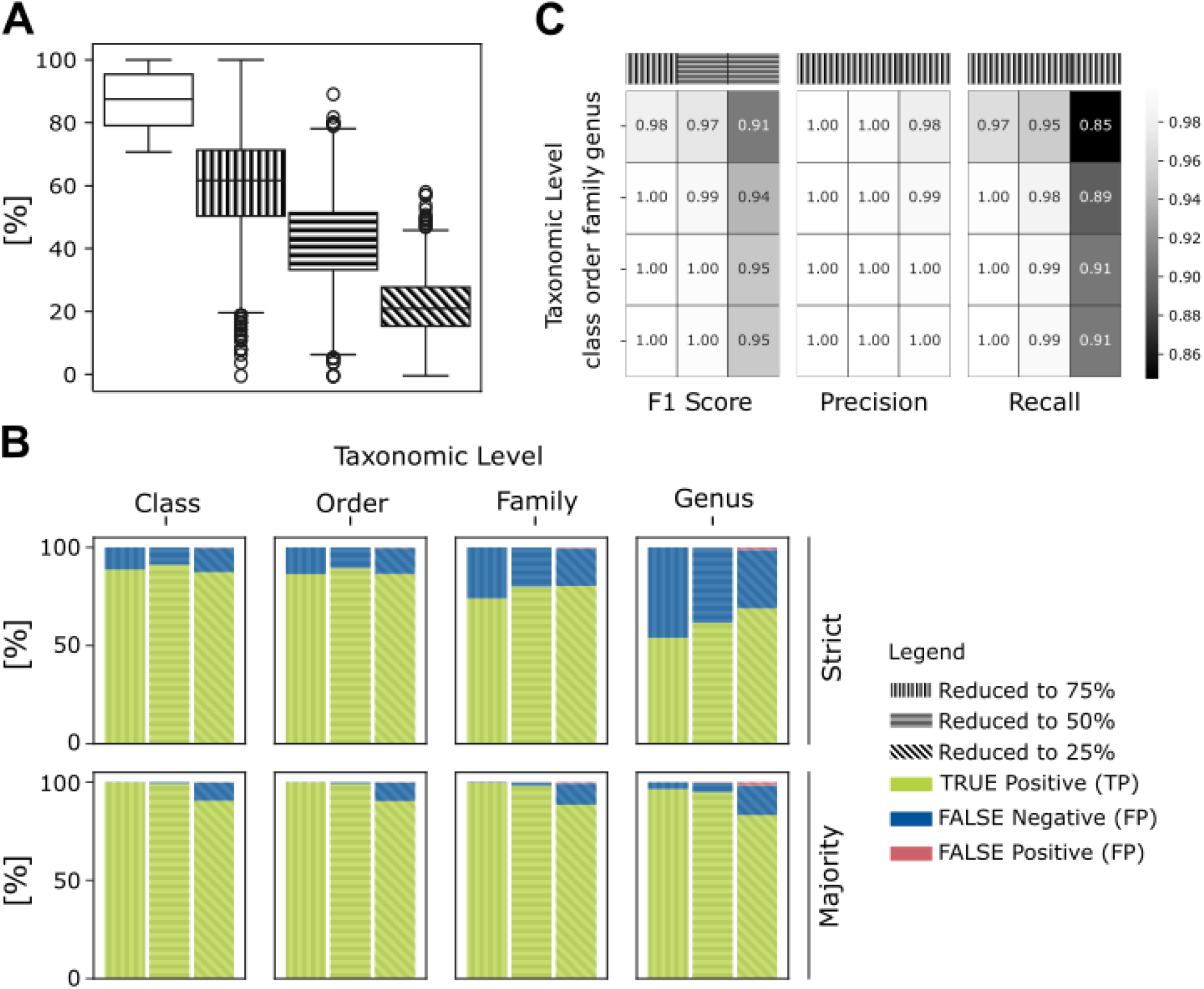
Taxonomic classification with GVClass. A) completeness percentage of all 1109 benchmarked GVs genomes before and after reduction to 75% 50% and 25% of their original length. B)Relative distribution of GVClass “strict” and “majority” taxonomic predictions of reduced GVs genomes after reduction C) Calculated precision of GVClass. Recall measures the proportion of actual positives correctly identified by the GVClass, while precision measures the proportion of positive predictions that are actually correct. The closer to 1 the better.

Using “majority” taxonomic assignments, GVClass demonstrated high efficiency across all ranks based on the percentage of true positives (TPs) at 94.96%, false negatives (FNs) at 4.69% and false positives at 0.35% of overall results regardless of the genome completeness (Fig.2B).

The highest proportion of TPs was observed at the Class-level, with a slight decrease at lower taxonomic ranks (Class 96.52% > Order 96.43% > Family 95.43% > Genus 91.46%). At each taxonomic rank, genomes reduced to 75% completeness showed a higher proportion of TPs at 98.93% and 0.06 FPs, which decreased as genome completeness was further reduced to 50% at 97.79% TPs and 0.23% FPs and 88.17% TPs and 0.76% FPs at 25% completeness. On average, regardless of the completeness and the taxonomic ranks, GVClass achieved 97% precision (instances predicted as positive that were actually positive) and 91% recall (positive instances successfully identified) (Fig.2C). These high values demonstrate GVClass’s reliability and efficiency correctly identifying giant virus sequences while minimizing false positives and false negatives.

Closer analysis revealed that GVClass showed both highest precision and recall in high taxonomic levels, with slight decreases at lower ranks (Fig. 2C). As expected, stronger variations in precision and recall were observed with reduced genome completeness (Fig. 2C). More challenging was correct identification at Genus level for highly incomplete genomes (25% completeness), with precision at 86%, and recall at 76%. This result underlines the importance of marker gene presence, which is typically directly linked to sufficient sequencing depth and assembly quality.

Given that a large proportion of our benchmark data consists of GVMAGs, we examined performance of GVClass on genomes of giant virus isolates. From a set of 121 giant virus isolate genomes affiliated with the orders *Algalvirales, Asfuvirales, Chitovirales, Imitevirales*, and *Pimascovirales* as well as the proposed *Pandoravirales* (Fig 3), 97% were correctly predicted up to the genus level, exception were *Clandestinovirus, Ectocarpus siliculosus virus* 1 and *Feldmannia species virus* (STable. 1). However, each of the 3 isolates represent the only representative of their respective genus, making it impossible for GVClass to correctly predict them as self-hits were excluded from the analysis.

**Fig 3:**
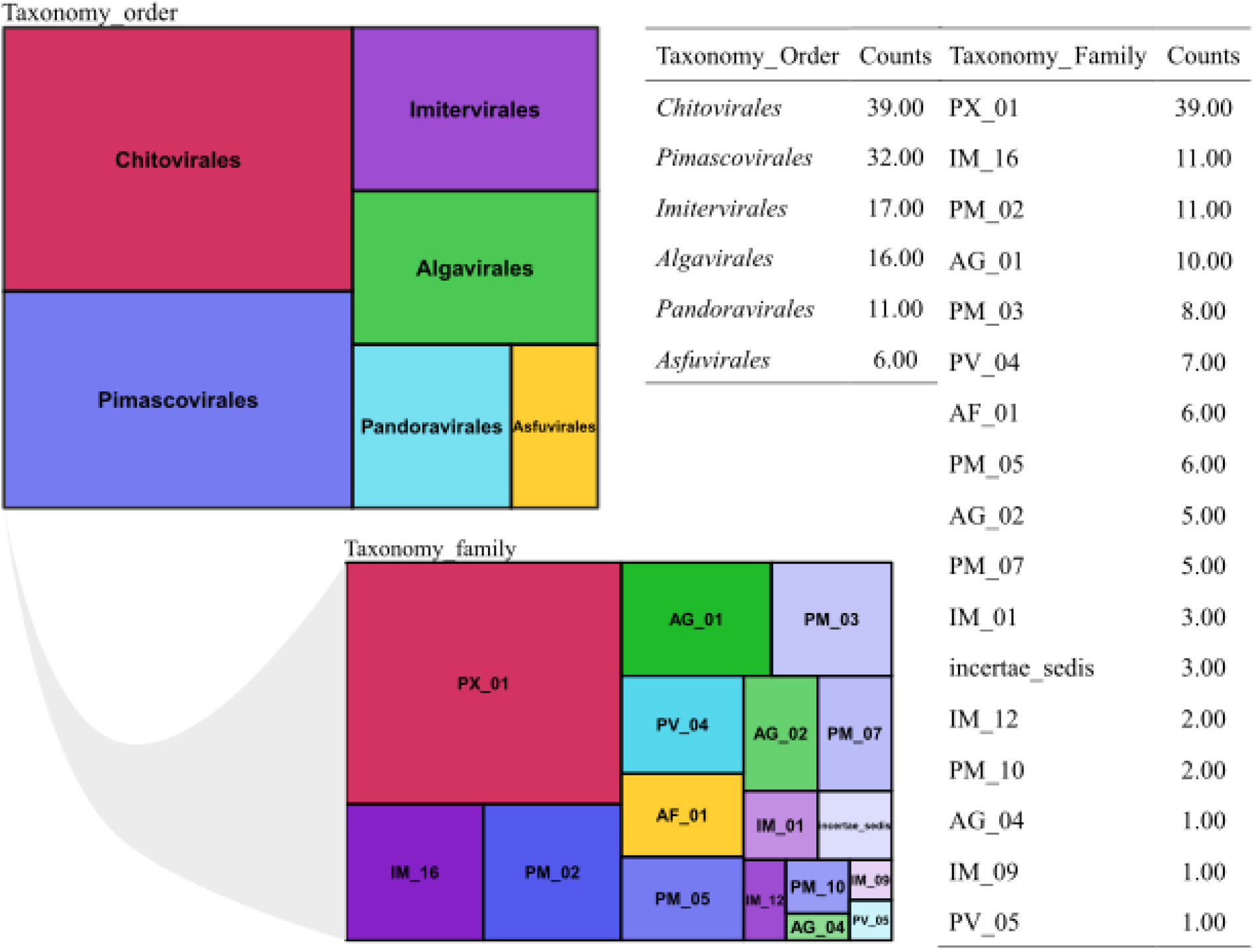
Treemap plot of the taxonomic classification of the 121 isolate genomes by GVClass

#### Completeness and Contamination Estimation with GVClass

Next, we tested the completeness estimation feature. We first ran GVClass on the original genomes to generate a baseline before screening the reduced genomes. The mean percentage of completeness of non-reduced genomes was estimated to 77.5% (Fig 2.A). Reduced genomes were respectively estimated to reach completeness of 61%, 42% and 22% respectively (Fig 2.A), closely matching the 75%, 50% and 25% reductions. With our set of isolates, the average completeness was estimated at 85.5% (Fig.4).

Based on our results we consider a completeness value below 30% as low, 30-70% as medium, and above 70% as high completeness when using GVClass. Our benchmarking illustrates the efficiency of GVClass in predicting giant virus query sequences with very high precision across taxonomic ranks and into estimating genome completeness.

In the second phase, we analyzed the order duplication factor of the tested isolates. Giant virus genomes typically have a duplication factor below 3, with higher values potentially indicating mixed viral populations. Based on our results an order duplication factor below 1.5 suggests a low chance of representing a mixed bin (high quality). An order duplication factor between 1.5 and 2 suggests a medium chance of a mixed bin (medium quality). Lastly, an order duplication factor above 3 suggests a high chance of a mixed bin (indicating low quality). Our analysis showed an average duplication factor of 1.3 confirming the high quality of our queries (Fig 4).

**Fig 4:**
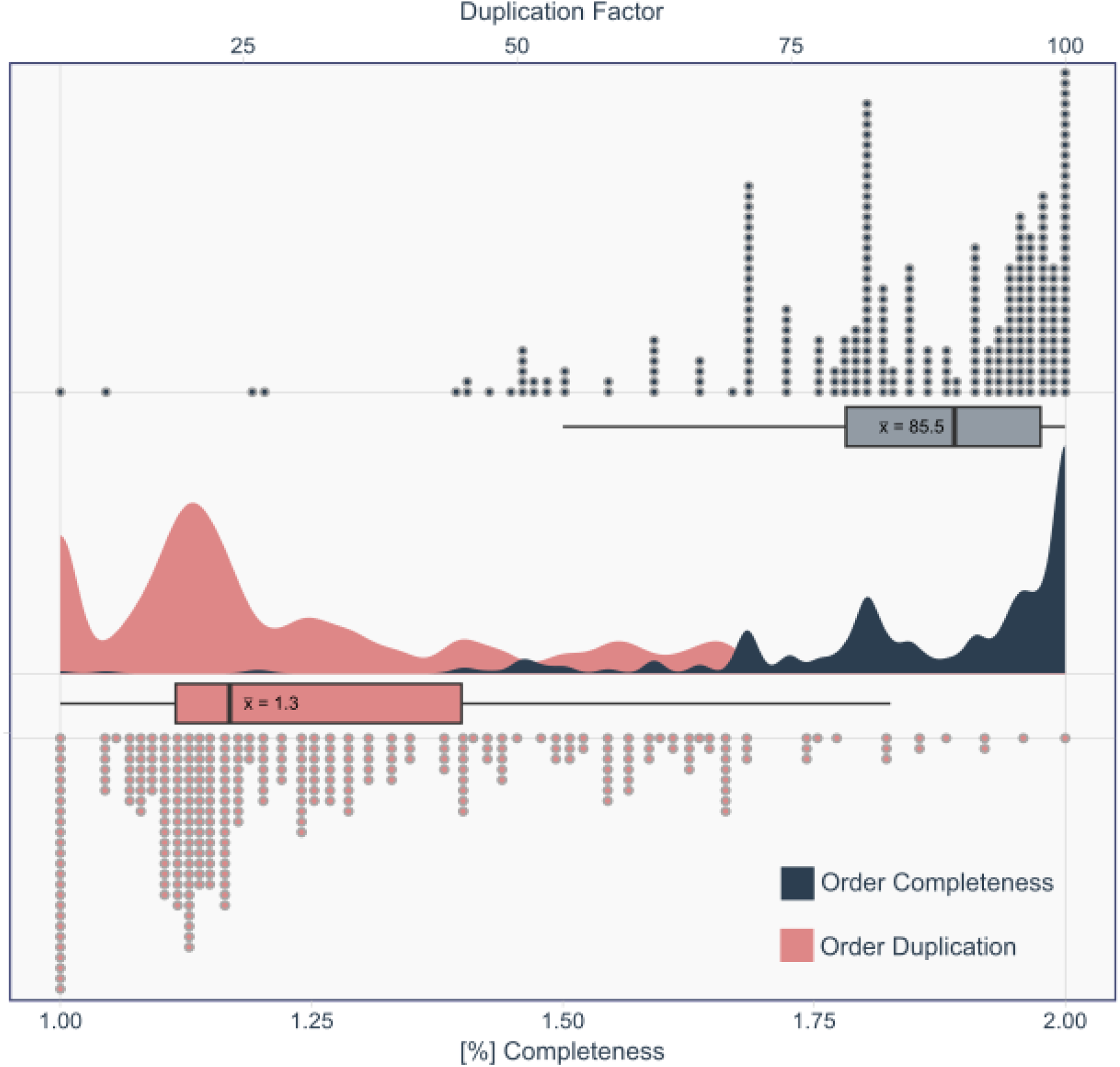
Distribution raincloud plot of duplication factor and percentage of completeness estimates by GVClass for the 121 isolates.

#### Novelty Detection Capability of GVClass

We tested GVClass’s capability to detect novelty using query sequences from 25 Nucleocytoviricota taxonomic groups (10 genera, 10 families, and 5 orders). To conduct this analysis, we created 25 versions of the database by removing all genomes associated with each selected group, one at a time. By running GVClass on the members of the omitted taxa, we could assess how the predictions perform on genomes that are novel at different taxonomic levels (Fig 5.A).

**Fig 5:**
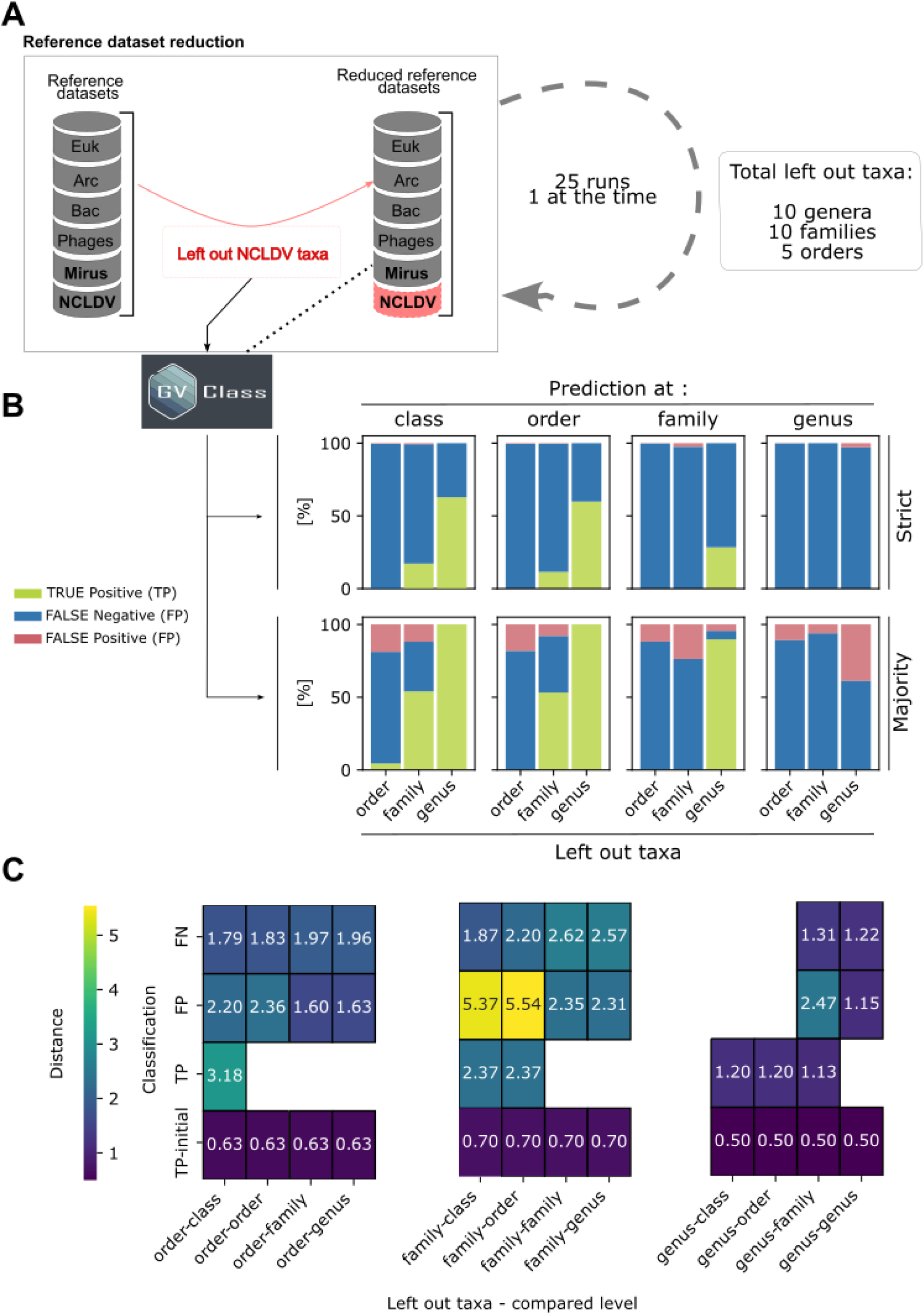
A) schematic of the reference dataset reduction for novelty detection. 25 versions of the database were created by removing all GVs genomes associated with each selected group, one at a time (10 genera, 10 families, and 5 orders). B) Relative distribution of GVClass “strict” and “majority” taxonomic predictions of omitted Nucleocytoviricota taxa C) Heatmap summarizing evolutionary distances to nearest neighbors for TP FP and FN for each group that was left out. For TP-initial (before genome reduction) self hits were excluded.

For instance, if a genus is removed from the database, members of that genus cannot be assigned to the correct genus, as it is no longer part of the reference data. This absence should result in a signal of novelty. Ideally, novelty should be detected as false negative (FN), as queries can not be assigned correctly due to missing data in the database.

Our results showed that the highest proportions of false negatives predictions were found at the same or higher taxonomic level than that of the omitted taxon (Fig 5.B). The main exception was for omitted genera, where classification at the genus level frequently yielded false positives. A possible explanation for this is the greater similarity between viruses of the same genus, which can lead to assignment to a closely related genus, resulting in false positives in the benchmarking (Fig 5.B).

We also estimated novelty by calculating the average phylogenetic distance for every predicted affiliation of omitted taxa. Our analysis revealed that the family level shows the highest novelty detection capability, indicated by the highest phylogenetic distances for false negatives. The order level has moderate novelty detection capability, and the genus level shows the least novelty for misclassified sequences (Fig 5.C). Considering both the proportion of predicted instances and the phylogenetic distance, we concluded that the family rank is the most suitable for novelty detection with GVClass.

Our benchmarking demonstrated GVClass’s robust capability to detect novel viral sequences, particularly at higher taxonomic ranks. This is crucial for identifying potentially new viral lineages in metagenomic data, which will add to the existing toolkit for exploring viral diversity and evolution.

## Acknowledgements

The work conducted by the U.S. Department of Energy Joint Genome Institute (https://ror.org/04xm1d337), a DOE Office of Science User Facility, is supported by the Office of Science of the U.S. Department of Energy operated under Contract No. DE-AC02-05CH11231.

## Code availability

GVClass is an open-source software, and its code can be found at https://github.com/NeLLi-team/gvclass.

## Competing interests

The authors declare that they have no competing interests.

## Notes

### Competing Interest Statement

The authors have declared no competing interest.

## Bibliography

Aylward, Frank O., and Mohammad Moniruzzaman. 2021. ‘Viralrecall—a Flexible Command-Line Tool for the Detection of Giant Virus Signatures in ‘omic Data’. Viruses 13 (2). 10.3390/v13020150.

Aylward, Frank O., Mohammad Moniruzzaman, Anh D. Ha, and Eugene V. Koonin. 2021. ‘A Phylogenomic Framework for Charting the Diversity and Evolution of Giant Viruses’. PLoS Biology 19 (10 October). 10.1371/JOURNAL.PBIO.3001430.

Camargo, Antonio Pedro, Stephen Nayfach, I. Min A. Chen, Krishnaveni Palaniappan, Anna Ratner, Ken Chu, Stephan J. Ritter, et al. 2023. ‘IMG/VR v4: An Expanded Database of Uncultivated Virus Genomes within a Framework of Extensive Functional, Taxonomic, and Ecological Metadata’. Nucleic Acids Research 51 (D1). 10.1093/nar/gkac1037.

Camargo, Antonio Pedro, Simon Roux, Frederik Schulz, Michal Babinski, Yan Xu, Bin Hu, Patrick S.G. Chain, Stephen Nayfach, and Nikos C. Kyrpides. 2023. ‘Identification of Mobile Genetic Elements with GeNomad’. Nature Biotechnology. 10.1038/s41587-023-01953-y.

Capella-Gutiérrez, Salvador, José M. Silla-Martínez, and Toni Gabaldón. 2009. ‘TrimAl: A Tool for Automated Alignment Trimming in Large-Scale Phylogenetic Analyses’. Bioinformatics 25 (15). 10.1093/bioinformatics/btp348.

Claverie, Jean Michel, and Chantal Abergel. 2016. ‘Giant Viruses: The Difficult Breaking of Multiple Epistemological Barriers’. Studies in History and Philosophy of Science Part C :Studies in History and Philosophy of Biological and Biomedical Sciences 59. 10.1016/j.shpsc.2016.02.015.

Filée, Jonathan, and Michael Chandler. 2010. ‘Gene Exchange and the Origin of Giant Viruses’. Intervirology. 10.1159/000312920.

Gaïa, Morgan, Lingjie Meng, Eric Pelletier, Patrick Forterre, Chiara Vanni, Antonio Fernandez-Guerra, Olivier Jaillon, et al. 2023. ‘Mirusviruses Link Herpesviruses to Giant Viruses’. Nature 616 (7958). 10.1038/s41586-023-05962-4.

Kerepesi, Csaba, and Vince Grolmusz. 2017. ‘The “Giant Virus Finder” Discovers an Abundance of Giant Viruses in the Antarctic Dry Valleys’. Archives of Virology 162 (6). 10.1007/s00705-017-3286-4.

Moniruzzaman, Mohammad, Carolina A. Martinez-Gutierrez, Alaina R. Weinheimer, and Frank O. Aylward. 2020. ‘Dynamic Genome Evolution and Complex Virocell Metabolism of Globally-Distributed Giant Viruses’. Nature Communications 11 (1): 1–12. 10.1038/s41467-020-15507-2.

Moniruzzaman, Mohammad, Alaina R Weinheimer, Carolina A Martinez-Gutierrez, and Frank O Aylward. 2020. ‘Widespread Endogenization of Giant Viruses Shapes Genomes of Green Algae.’ Nature 588 (7836): 141–45. 10.1038/s41586-020-2924-2.

Natalya, Yutin, Mutz Pascal, Krupovic Mart, and Koonin Eugene V. 2024. ‘Mriyaviruses: Small Relatives of Giant Viruses’. MBio 15 (7): e01035–24. 10.1128/mbio.01035-24.

Nguyen, Lam Tung, Heiko A. Schmidt, Arndt Von Haeseler, and Bui Quang Minh. 2015. ‘IQ-TREE: A Fast and Effective Stochastic Algorithm for Estimating Maximum-Likelihood Phylogenies’. Molecular Biology and Evolution 32 (1). 10.1093/molbev/msu300.

Parks, Donovan H., Maria Chuvochina, Christian Rinke, Aaron J. Mussig, Pierre Alain Chaumeil, and Philip Hugenholtz. 2022. ‘GTDB: An Ongoing Census of Bacterial and Archaeal Diversity through a Phylogenetically Consistent, Rank Normalized and Complete Genome-Based Taxonomy’. Nucleic Acids Research 50 (D1). 10.1093/nar/gkab776.

Pitot, Thomas M, Josephine Z Rapp, Frederik Schulz, Catherine Girard, Simon Roux, and Alexander I Culley. 2024. ‘Distinct and Rich Assemblages of Giant Viruses in Arctic and Antarctic Lakes’. ISME Communications, January. 10.1093/ismeco/ycae048.

Richter, Daniel J., Cédric Berney, Jürgen F. H. Strassert, Yu-Ping Poh, Emily K. Herman, Sergio A. Muñoz-Gómez, Jeremy G. Wideman, Fabien Burki, and Colomban de Vargas. 2022. ‘EukProt: A Database of Genome-Scale Predicted Proteins across the Diversity of Eukaryotes’. Peer Community Journal 2. 10.24072/pcjournal.173.

Schulz, Frederik, Chantal Abergel, and Tanja Woyke. 2022. ‘Giant Virus Biology and Diversity in the Era of Genome-Resolved Metagenomics’. Nature Reviews Microbiology. Nature Research. 10.1038/s41579-022-00754-5.

Schulz, Frederik, Julien Andreani, Rania Francis, Jacques Yaacoub Bou Khalil, Janey Lee, Bernard La Scola, and Tanja Woyke. 2020. ‘Advantages and Limits of Metagenomic Assembly and Binning of a Giant Virus’. BioRxiv 5 (3): 1–10. 10.1101/2020.01.10.902254.

Schulz, Frederik, Simon Roux, David Paez-Espino, Sean Jungbluth, David A. Walsh, Vincent J. Denef, Katherine D. McMahon, et al. 2020. ‘Giant Virus Diversity and Host Interactions through Global Metagenomics’. Nature 578 (7795): 432–36. 10.1038/s41586-020-1957-x.

Schulz, Frederik, Natalya Yutin, Natalia N Ivanova, Davi R Ortega, Tae Kwon Lee, Julia Vierheilig, Holger Daims, et al. 2017. ‘Giant Viruses with an Expanded Complement of Translation System Components’. Science 85 (April): 82–85.

Scola, Bernard La, Stéphane Audic, Catherine Robert, Liang Jungang, Xavier De Lamballerie, Michel Drancourt, Richard Birtles, Jean Michel Claverie, and Didier Raoult. 2003. ‘A Giant Virus in Amoebae’. Science 299 (5615): 2033. 10.1126/science.1081867.

Verneau, Jonathan, Anthony Levasseur, Didier Raoult, Bernard La Scola, and Philippe Colson. 2016. ‘MG-Digger: An Automated Pipeline to Search for Giant Virus-Related Sequences in Metagenomes’. Frontiers in Microbiology 7 (MAR). 10.3389/fmicb.2016.00428.

